# Rationally Designed PKD1 Activator Protects Against Neurodegeneration in Pre-clinical Models of Parkinson’s Disease

**DOI:** 10.1101/2025.02.13.637661

**Authors:** Arunkumar Asaithambi, Ahyoung Jang, Anamitra Ghosh, Muhammet Ay, Huajun Jin, Vellareddy Anantharam, Arthi Kanthasamy, Anumantha G. Kanthasamy

## Abstract

Oxidative stress leads to degeneration in Parkinson’s disease (PD). The key signal transduction and regulatory networks that are involved during this degenerative process in PD are currently being investigated for novel neuro-protective strategies. We recently discovered that the activation of Protein Kinase D1 (PKD1) acts as a novel compensatory mechanism in PD models and positive modulation of PKD1 can be a therapeutic strategy. Therefore, the purpose of the present study was to take a translational approach by developing a PKD1 activator and characterizing the protective function in pre-clinical models of PD. Positive genetic modulation of PKD1 by overexpression of constitutively active PKD1 protected against MPP^+^ induced dopaminergic neurotoxicity. Pharmacological activation by Rosiglitazone protected, whereas inhibition by kb NB 142-70 exacerbated against MPP^+^ and 6-OHDA toxicity in cell culture PD models. Importantly, peptides were rationally designed and screened for their ability to activate PKD1 using our screening methods. Peptide AK-P4 was identified to activate PKD1 specifically and protect against MPP^+^ and 6-OHDA in both N27 cells and primary mesencephalic neurons. Further AK-P4 tagged with TAT sequence (AK-P4T) delivered using intra-venous injections activated PKD1 in mice. The neuro-protective effects of AK-P4T were tested using the sub-chronic MPTP mice model. Co-treatment with AK-P4T significantly restored the neurotransmitter levels and the behavioral and locomotory activities of the MPTP mouse model of PD. Collectively, our results demonstrate that rationally designed PKD1 activator peptide AK-P4T positively modulated PKD1 and protected against neurodegeneration in the pre-clinical models of PD. Our results suggest that positive modulation of the PKD1 using AK-P4T shows promise as a potential therapeutic agent against PD.

## Introduction

Parkinson’s disease (PD) is the second common neurodegenerative disorder affecting over a million Americans. Current treatment approaches available for Parkinson’s disease are symptomatic and fail to prevent the progression of the neurodegenerative process. The currently available drugs are limited in their effectiveness to either slow or stop the progressive neurodegenerative processes in PD, largely due to the lack of mechanistic insights into the selective dopaminergic degenerative process [1–3].

Experimental findings from cell cultures, animal models, and humans indicate that oxidative stress and apoptosis may play a major role in the pathophysiological processes underlying PD. Neuronal cells maintain an oxidant/ antioxidant homeostatic balance. Any breach in this homeostasis causes excessive ROS production and oxidative damage, which might lead to neurodegenerative disorders [1, 2, 4–8].

Most investigations of signaling in neurodegenerative disorders including ours have primarily focused on signaling pathways that is involved in execution of cell death during this oxidant / antioxidant homeostatic breach [9–13]. None of the therapeutic interventions developed against the major cell death kinases have progressed to the treatment stage, probably due to the lack of understanding of the complex signal transduction process [14]. This resulted in an imperative to find an alternative approach to prevent PD progression.

The protective compensatory mechanisms to counteract the oxidative damage in Parkinson’s disease are still largely unexplored. Recently, we reported that protein kinase D1 (PKD1) an oxidative stress sensor is activated during the early stages of oxidative stress to maintain the homeostatic balance and promote cell survival in PD cell culture and animal models [15]. PKD1 get fully activated through phosphorylation at activation loop serine sites (S744/S748). Further phosphorylation at PKD1 S916 proceeds and regulates the activation loop phosphorylation [15].

We hypothesized that positive modulation of PKD1 can be a novel therapeutic approach against PD. So, we rationally designed peptide modulators to activate PKD1. The PKD1 activator peptide AK-P4T protected against neurodegeneration in both cell culture and animal models of PD. Herein, we describe the design, development, and characterization of the peptide-based drug AK-P4T as a novel therapeutic agent in pre-clinical models of Parkinson’s disease.

## Materials and Methods

### Cell Culture

The immortalized rat mesencephalic dopaminergic neuronal cell line (N27) was a kind gift from Dr. Kedar N. Prasad (University of Colorado Health Sciences Center, Denver, CO). N27 cells were grown in RPMI 1640 medium containing 10% fetal bovine serum, 2 mm L-glutamine, 50 units of penicillin, and 50 μg/ml streptomycin. Cells were maintained in a humidified atmosphere of 5% CO_2_ at 37 °C, as described previously [16]. N27 cells are used widely as a cell culture model for PD [13, 16–20].

### Primary mesencephalic neuronal culture

Primary mesencephalic neuronal cultures were prepared from the ventral mesencephalon of gestational 16- to 18-day-old mouse embryos, as described earlier [21]. Tissues were dissected from E16 to E18 mouse embryos maintained in ice cold Ca2^+^-free Hanks’ balanced salt solution and then dissociated in Hanks’ balanced salt solution containing trypsin-0.25% EDTA for 30 min at 37°C. The dissociated cells were then plated at equal density (0.5 × 10^6^ cells) on 12 mm coverslips precoated with 0.1 mg/ml poly-D-lysine. Cultures were maintained in neurobasal medium fortified with B- 27 supplements, 500 μM l-glutamine, 100 IU/ml penicillin, and 100 μg/ml streptomycin (Invitrogen). The cells were maintained in a humidified CO_2_ incubator (5% CO_2_ and 37°C). Half of the culture medium was replaced every 2 days. Approximately 6- to 7-day-old cultures were used for experiments. Primary mesencephalic dopaminergic neuronal cells were exposed to 10 μM for 1 h.

### Cytotoxicity Assays

Cell death was determined using the Sytox green cytotoxicity assay, after exposing the N27 cells to H_2_O_2_ (100 μm), as described previously. This cytotoxicity assay was optimized for a multiwell format, which is more efficient and sensitive than other cytotoxicity measurements [22, 23]. Briefly, N27 cells were grown in 24-well cell culture plates at 100,000 cells per well and treated with H_2_O_2_ (100 μm) and 1 μm Sytox Green fluorescent dye. The Sytox green assay allows dead cells to be viewed directly under a fluorescence microscope, as well as quantitatively measured with a fluorescence microplate reader (excitation 485 nm; emission 538 nm) (Biotek). Phase contrast and fluorescent images were taken after H_2_O_2_ exposure with a NIKON TE2000 microscope, and pictures were captured with a SPOT digital camera.

### Immunocytochemistry

The primary mesencephalic neurons or N27 cells after H_2_O_2_ treatment were fixed with 4% paraformaldehyde and processed for immunocytochemical staining. First, nonspecific sites were blocked with 2% bovine serum albumin, 0.5% Triton and 0.05% Tween-20 in phosphate-buffered saline (PBS) for 20 min. The cells then were incubated with antibodies directed against TH, native PKD1 and PKD1-pS744/S748 in PBS containing 1% BSA at 4°C overnight, followed by incubation with Alexa 488 and Alexa 568 conjugated secondary antibodies in PBS containing 1% BSA. Secondary antibody treatments were followed by incubation with Hoechst 33342 dye for 5 min at room temperature to stain the nucleus. The coverslips containing stained cells were washed with PBS, mounted on slides, and viewed under a Nikon inverted fluorescence microscope (model TE-2000U; Nikon, Tokyo, Japan). Both fluorescence and confocal images were captured with a SPOT digital camera (Diagnostic Instruments, Inc., Sterling Heights, MI).

### Immunohistochemical staining of brain slices

Mice were perfused with 4% paraformaldehyde and then the brain slices were cut on a microtome into 20 μm sections. Brain sections were blocked with PBS containing 1% BSA and 0.1% Triton X-100 in for 20 min and then incubated with antibodies directed against PKD1, PKD1- pS744/S748 and TH overnight at 4°C followed by incubation with either Alexa 488-conjugated (green, 1:1,000) or Alexa 568-conjugated (red, 1:1,000) secondary antibody for 1 h at RT. Secondary antibody treatments were followed by incubation with Hoechst 33342 (10 μg/ml) for 3 min at RT to stain the nucleus. Then the slices were mounted on a slide and viewed under a Nikon inverted fluorescence microscope (model TE-2000U); images were captured with a SPOT digital camera (Diagnostic Instruments, Sterling Heights, MI).

### Diaminobenzidine immunostaining and stereological counting of TH- positive neurons

Tyrosine hydroxylase–diaminobenzidine (DAB) immunostaining was performed in striatal and substantia nigral sections as described previously. Mice brains after sacrifice were perfused with 4% paraformaldehyde (PFA) and postfixed with PFA and 30% sucrose. The fixed brains were subsequently cut using previous termanext term cryostat into 30-μm coronal sections and kept in 30% sucrose–ethylene glycol solution at −20 °C. On the day of staining, sections were rinsed with PBS and incubated with the anti-TH antibody (Calbiochem rabbit anti-mouse, 1:1,600) overnight at 4 °C. Biotinylated anti-rabbit secondary antibody was used for 1 h at room temperature. The sections were then incubated with avidin peroxidase (Vectastatin ABC Elite kit) for 30 min at room temperature. Immunolabeling was observed using DAB, which yielded a brown stain. Total numbers of TH-stained neurons in SNpc were counted stereologically with Stereo Investigator software (MicroBrightField, Williston, VT, USA), using an optical fractionator.

### Western Blot Analysis

Cells were lysed in either modified RIPA buffer or M-PER buffer (Thermo Scientific) for Western blot, immunoprecipitation and kinase assays. Lysates containing equal amounts of protein were loaded in each lane and separated on 10-12% SDS-PAGE, as described previously (Kaul et al., 2003). PKD1 polyclonal (1:1,000), PKCδ polyclonal (1: 1,000), PKD1-pS744/S748 (1:1,000), PKD1-pS916 (1:1,000), PKD1-pY469 (1:1,000), -and β-actin (1:10,000) antibodies were used to blot the membranes. IR dye- 800 conjugated anti-rabbit (1:5,000) and Alexa Fluor 680 conjugated anti-mouse (1:10,000) were used for antibody detection with the Odyssey IR Imaging system (LICOR), as previously described.

### PKCδ Kinase Assay

Immunoprecipitation and PKCδ kinase assay were performed as described earlier [13]. After cell lysis, cells were immunoprecipitated using a polyclonal PKCδ rabbit antibody and protein A Sepharose, and washed three times with PKCδ kinase buffer (40 mM Tris (pH 7.4), 20 mM MgCl_2_, 20 μM ATP, 2.5 mM CaCl2). The reaction was started by adding 20 μl of buffer containing 0.4 mg histone and 5 μCi of [γ-^32^P]ATP (4,500 Ci/mM).

After incubation for 10 min at 30°C, SDS loading buffer (2X) was added to the samples to terminate the reaction. The reaction products were separated on SDS-PAGE (12%), and the H1-phosphorylated bands were detected using a phosphoimager (Fujifilm FLA-5100) and quantified with MultiGauge V3.0 software.

### Protein Kinase D1 Kinase Assay

The cells were exposed to H_2_O_2_ (100 μM) for 1 h and cell lysates were immunoprecipitated, as previously reported, with native PKD1 antibody (Santa Cruz). The kinase reaction was carried out at room temperature for 20 min after adding 10 μl of kinase substrate mix (0.1 mM ATP + 10 µci [γ-^32^P] ATP + 2 μg Syntide 2 peptide substrate in kinase buffer). Kinase buffer contains 20 mM Tris pH 7.5, 10 mM MgCl_2_, and 1 mM DTT. Kinase assay performed using recombinant PKD1 protein in cell free system contained 90 μg/ml Phosphatidyl serine and 175 nM PMA in addition in the assay buffer. The samples were centrifuged to terminate the kinase reaction, and the supernatants containing the phosphorylated peptide were applied as spots to P81 phosphocellulose squares (Whatmann). The papers were washed four times with 0.75% phosphoric acid and once with acetone and dried, and activity was determined by liquid scintillation counting. The samples were also loaded on a SDS-PAGE and probed for native PKD1 to determine equal loading.

### Transient and Stable Transfections

PKD1 activation loop, active PKD1S744E/S748E (PKD1-CA) were obtained from Addgene, Inc. [24]. Electroporation was carried out with an Amaxa Nucleofector instrument, as per the manufacturer’s protocol. The transfected cells were then transferred to T-75 flasks or 6-well plates as desired and allowed to grow for a 24 h period before the treatment.

### Uptake of [^3^H] Dopamine

The effects of PKD1 inhibitor AK-P4T on the uptake of dopamine were assessed in fetal mouse mesencephalic cultures using [^3^H]dopamine (DA), as described previously (Afeseh et al., 2009). In brief, after incubation for 48 h with 5 μM MPP^+^, with or without kb NB 142-70, medium with the treatment was removed and cells were then washed once by assay incubation (Krebs-Ringer) buffer (5.6 mM glucose, 1.3 mM EDTA, 1.2 mM magnesium sulfate, 1.8 mM calcium chloride, 4.7 mM potassium chloride, 120 mM sodium chloride, and 16 mM sodium phosphate). Cells were incubated with 10 μM [^3^H] DA (30 Ci/mol) for 20 min at 37°C. Positive controls were obtained by incubating the cells with 10 μM [^3^H] DA together with 1 nM mazindol (potent dopamine reuptake inhibitor). The uptake was stopped by removing the reaction mixture and followed by three washes with fresh Krebs-Ringer buffer. Cells were then collected with the use of 1 N NaOH, and the radioactivity was measured by liquid scintillation counting.

### HPLC analysis of striatal dopamine and its metabolite levels

High-performance liquid chromatography (HPLC) with electrochemical detection were used to quantify striatal dopamine (DA), 3,4- dihydroxyphenylacetic acid (DOPAC), and homovanillic acid (HVA) levels. Samples Briefly, after 7 days of MPTP injection, mice were sacrificed and striata were collected and stored at − 80 °C. On the day of analysis, neurotransmitters from striatal tissues were extracted in 0.1 M perchloric acid solutions containing 0.05% Na2EDTA and 0.1% Na2S2O5 and isoproterenol (as an internal standard). The extracts were filtered in 0.22- μm spin tubes, and 20 μl of the sample was loaded for analysis. DA, DOPAC, and HVA were separated isocratically in a reversed-phase column using a flow rate of 0.7 ml/min. An HPLC system (ESA, Bedford, MA, USA) with an automatic sampler equipped with a refrigerated temperature control (Model 542; ESA) was used in these experiments. The electrochemical detection system consisted of a Coulochem Model 5100A with a microanalysis cell (Model 5014A) and a guard cell (Model 5020; ESA). Standard stock solutions of catecholamines (1 mg/ml) were prepared in perchloric acid solution and further diluted to a final working concentration of 50 pg/μl before injection. The data acquisition and analysis were performed using the EZStart HPLC software (ESA).

### Animals and treatment

Six- to 8-week-old male C57BL/6 mice weighing 24 to 28 g were housed under standard conditions: constant temperature (22 ± 1 °C) and humidity (relative, 30%) and a 12-h light/dark cycle. Mice were allowed free access to food and water. Use of the animals and protocol procedures were approved and supervised by the Committee on Animal Care at Iowa State University (Ames, IA, USA). Mice received AK-P4T (5 mg/kg dose) and the control peptide intravenous injections for 1 day (pretreatment before MPTP administration), 5 days (co-treatment with MPTP), and 7 days (post-treatment with MPTP). In the subchronic MPTP regimens, MPTP (25 mg/kg) was injected intraperitoneally via a single dose daily starting on day 2 for 5 days. The control mice received saline at the same dose.

### Behavioral measurements

An open-field experiment for testing locomotor activities after MPTP and AK-P4T treatments An automated device (AccuScan, Model RXYZCM- 16; Columbus, OH, USA) was used to measure the spontaneous activity of the mice. The activity chamber was 40 × 40 × 30.5 cm, made of clear Plexiglas, and covered with a Plexiglas lid with holes for ventilation. The infrared monitoring sensors were located every 2.54 cm along the perimeter (16 infrared beams along each side) and 2.5 cm above the floor. Two additional sets of 16 sensors were located 8.0 cm above the floor on opposite sides. Data were collected and analyzed by a VersaMax analyzer (AccuScan, Model CDA-8). Before any treatment, mice were placed inside the infrared monitor for 10 min daily for 3 consecutive days to train them. Five days after the last MPTP injection, both open-field and rotarod experiments were conducted.

### Statistical Analysis

Data analysis was performed using Prism 3.0 software (GraphPad Software, San Diego, CA). Bonferroni’s multiple comparison testing was used to find the significant differences between treatment and control groups. Differences with p < 0.05, p < 0.01, and p < 0.001 were considered significantly different from n≥6 from two or more independent experiments, and are indicated in the figures.

## Results

### Genetic and Pharmacological modulation of PKD1 activation protects against cell death in PD cell culture model

We used the two major parkinsonian toxicants 6-OHDA and MPP^+^ in the current study. We have previously shown that PKD1 is activated in 6- OHDA and MPP^+^ induced oxidative stress [15]. As shown in Fig. 1A-B, 100 µM 6-OHDA and 300µM MPP^+^ induces PKD1 activation loop phosphorylation (pS744/pS748) in a time dependent manner in N27 dopaminergic cells. Further, positive and negative genetic manipulation of PKD1 confirms the cell survival role of PKD1. We overexpressed the constitutively active human PKD1 plasmid (PKD1^-S744E/S748E^) in N27 cells and then exposed to 6-OHDA and MPP^+^. Mutation of serine residues at position 744 and 748 results in constitutively active PKD1 [24]. As shown in Fig. 1C, overexpression of PKD1^S744E^/^S748E^ significantly attenuated 6- OHDA and MPP^+^ induced cytotoxic cell death compared to vector transfected dopaminergic cells.

**Figure 1.**
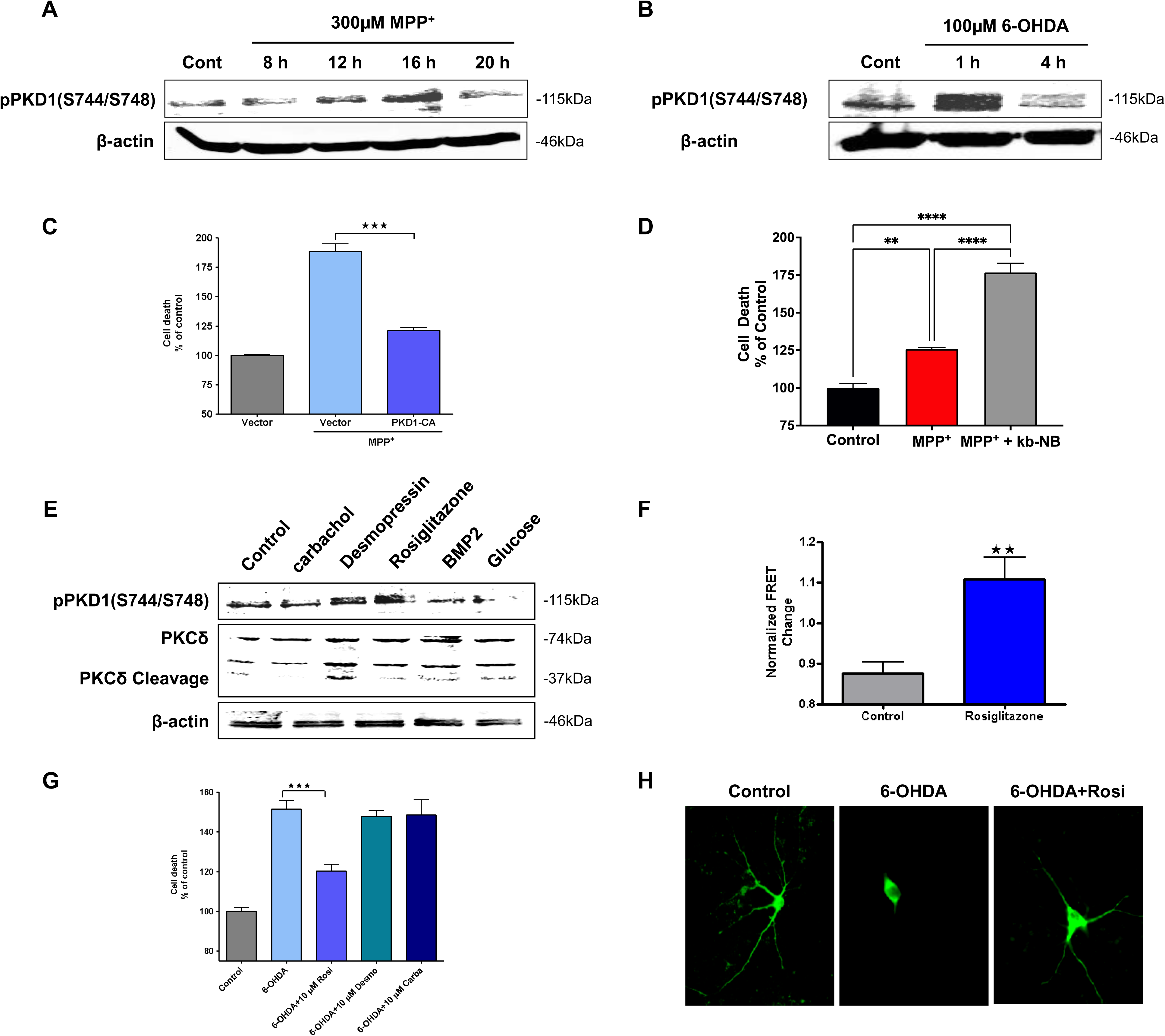
Pharmacological activation of PKD1 by Rosiglitazone shows neuroprotection in N27 cells and primary neurons. N27 dopaminergic cells were treated with MPP^+^ (300 µM) for 0-20 h and 6-OHDA (100 µM) for 0-4 h and blotted for PKD1 activation loop phosphorylation pS744/pS748 (A & B). N27 dopaminergic cells transiently transfected with 5 μM PKD1 ^S744E/S748E^ and 5 μM vector plasmid were treated with or without 300 μM MPP^+^ and monitored for cytotoxicity at various time points using Sytox green. (C). N27 cells were co-treated with or without 50 µM PKD1 inhibitor kb NB 14270 and monitored for cell death during MPP^+^ treatment (D). N27 cells were treated with 10 µM Carbachol, 50 µM Desmopressin, 100 µM Rosiglitazone, 100 ng/ml BMP2, and 25mM Glucose and probed for PKD1 (pS744E/pS748E) and PKCδ cleavage (E). N27 cells transfected with 5μM PKD1 kinase activity reporter (PKD1-KAR) plasmid and 5 μM vector plasmid were treated with Rosiglitazone and monitored for CFP/YFP FRET change in a high-throughput format (F). N27 cells were co-treated with or without 10 µM Rosiglitazone, 10 µM Carbachol, or 10 µM Desmopressin and monitored for cell death during 6-OHDA treatment (G). Tyrosine Hydroxylase (TH) positive primary mesencephalic neurons treated with co-treated with or without 10 µM Rosiglitazone were monitored for changes in neuronal morphology and neuroprotection using immune-fluorescence (H). **p<0.01 and ***p<0.001 denote significant differences between groups from n≥6. (n=8).

Since genetic modulation of PKD1 can protect neuronal cells, we hypothesized that pharmacological modulation of PKD1 should influence cell survival and death. N27 dopaminergic cells were exposed to 300 μM MPP^+^ in the presence or absence of 50 μM PKD inhibitor kb NB 142-70 and cytotoxic cell death was determined using Sytox green assay. As shown in Fig. 1D, co-treatment with 50 μM kb NB 142-70 exacerbated MPP^+^-induced cytotoxic cell death compared to MPP^+^ alone treated cells.

Next, as a proof of concept we examined if positive pharmacological modulation of PKD1 can protect against oxidative damage and cell death. Since there are no PKD1 activators available, we screened for PKD1 activators. We have previously shown that in dopaminergic neurons proteolytically activated PKCδ phosphorylates PKD1 during the early stage, so we looked for drugs that can cause persistent activation of PKD1 independent of PKCδ cleavage. After carefully reviewing different PKD1 activation mechanisms in various other models, we handpicked some drugs and screened them for their potential to activate PKD1 through these mechanisms. As shown in Fig. 1E, primary screening of 10 µM Carbachol, 50 µM Desmopressin, 100 µM Rosiglitazone, 100 ng/ml BMP2, and 25mM Glucose were performed using western blot to screen for their potential to phosphorylate activation loop of PKD1 (pS744E/pS748E). Even though both Rosiglitazone and Desmopressin induced PKD1 activation loop phosphorylation, only Rosiglitazone’s mechanism of activation was independent of PKCδ cleavage. Next, we used a genetic based FRET reporter optimized in a plate reader format as a secondary screening method to report PKD1 activity. Previously, this FRET reporter has been used to monitor PKD1 activity directly in cells [25–27]. As shown in Fig. 1F, 10 µM Rosiglitazone increased PKD1 activity as shown by the FRET change (CFP/YFP) read out. Further, only 10 µM Rosiglitazone protected against 100 µM 6-OHDA induced cell death in N27 cells (Fig. 1G). Next, we examined if Rosiglitazone could protect against mouse primary dopaminergic neurons. As shown in Fig. 1H, Immunofluorescence pictures of Tyrosine Hydroxylase (TH) positive dopaminergic neurons showed protection against 10 µM 6-OHDA when co-treated with 10 µM Rosiglitazone. These results unequivocally demonstrate that positive pharmacological modulation of PKD1 has protective compensatory role against oxidative damage.

### Rational drug design

Even though we showed that positive modulation of PKD1 through Rosiglitazone has protective function in cell culture models of PD. We wanted to develop specific activators of PKD1, as rosiglitazone is not specific and has other known functions [25, 26]. Previously, it has been shown that specific peptide-based regulators targeting protein-protein interactions of PKCs have been successfully developed and advanced to clinical trials in cardiovascular diseases [28, 29]. PKD1 has a very different structure compared to PKCs; it has two cysteine rich domains and a pleckstrin homology (PH) domain in the regulatory fragment. It was shown that the regulatory fragment importantly the PH domain has an auto- inhibitory role by interacting with the catalytic fragment but lacks a defined pseudo-substrate domain like PKCs [30, 31]. After careful analysis of PKD1 primary and secondary structure, we hypothesized that PKD1 region containing the consensus substrate-like motif (LXRXXS/T) and pseudosubstrate-like motif (LXRXX#, LXR##, #- serine or threonine residue replaced mostly by a hydrophobic/aliphatic residue) can regulate PKD1 intra-protein interactions. Peptides mimicking these regions can be highly specific modulators of PKD1 activity. As shown in Fig. 2A, alignment of distantly related PKD1 sequences from murine, rat and C. elegans show the presence of highly conserved regions around the substrate-like and pseudosubstrate-like sequences only in the regulatory fragment. Further, the regions containing the pseudo-substrate like sequence are highly conserved between the three PKD isoforms, whereas the substrate-like sequence is present only in PKD1 (Fig. 2B). We chose four of these regions from each domain; the highly conserved 6-aa peptide AK-P1 (6-11 amino acids), and 11-aa peptide AK-P2 (36-46 amino acids) were synthesized from the cysteine rich domain 1, 10-aa peptide AK-P3 (295-395 amino acids) were synthesized from the cysteine rich domain 2, 12-aa peptide AK- P4 (405-416 amino acids) were synthesized from the pleckstrin homology (PH) domain. Additionally, peptides disrupting protein-protein interactions have also been shown to regulate kinase activity. Mochly-Rosen group reported that there are regions in PKC enzymes that are homologous to the PKC carrier protein RACK (termed pseudo-RACKS) and the peptides synthesized mimicking this region can facilitate PKC translocation and activation [32]. Using this rational, we detected similar regions in PKD1 that are homologous to PKD1 interaction proteins 14-3-3γ and Gβγ. As shown in Fig. 2C, a 5-aa peptide AK-P5 (54-58 amino acids) from the cysteine rich domain 1 (pseudo-14-3-3γ), a 9-aa peptide AK-P6 (79-87 amino acids) from the pleckstrin homology (PH) domain (pseudo-Gβγ) and a 5-aa control peptide AK-P7 (135-139 amino acids) were synthesized. Primary screening of the synthesized peptides was done using the phospho-specific antibodies to detect PKD1 S744/S748 and S916 phosphorylation. As shown in Fig. 2D, 50 µM of AK-P3 and AK-P4 induced PKD1 S744/S748 and S916 phosphorylation in N27 cells. Secondary screening was performed by monitoring PKD1 kinase activity as measured by a [^32^P] kinase assay using Syntide 2 substrate (Fig. 2E). Peptides AK-P3 and AK-P4 significantly increased PKD1 activity validating the results obtained from primary screening, whereas AK-P1 significantly decreased kinase activity. The specificity of the peptides in activating PKD1 was tested in a cell-free system using a human recombinant PKD1 enzyme. As shown in Fig. 2F, AK-P4 increased PKD1 kinase activity and directly activates PKD1. Finally, the genetic-based FRET reporter assay confirmed that AK-P4 treatment enhanced PKD1 activity in N27 cells (Fig. 2G). Together, these results confirm that peptides mimicking PKD1’s substrate-like region acts an inhibitor whereas peptides mimicking highly conserved PKD1’s pseudosubstrate-like region acts an activator.

**Figure 2.**
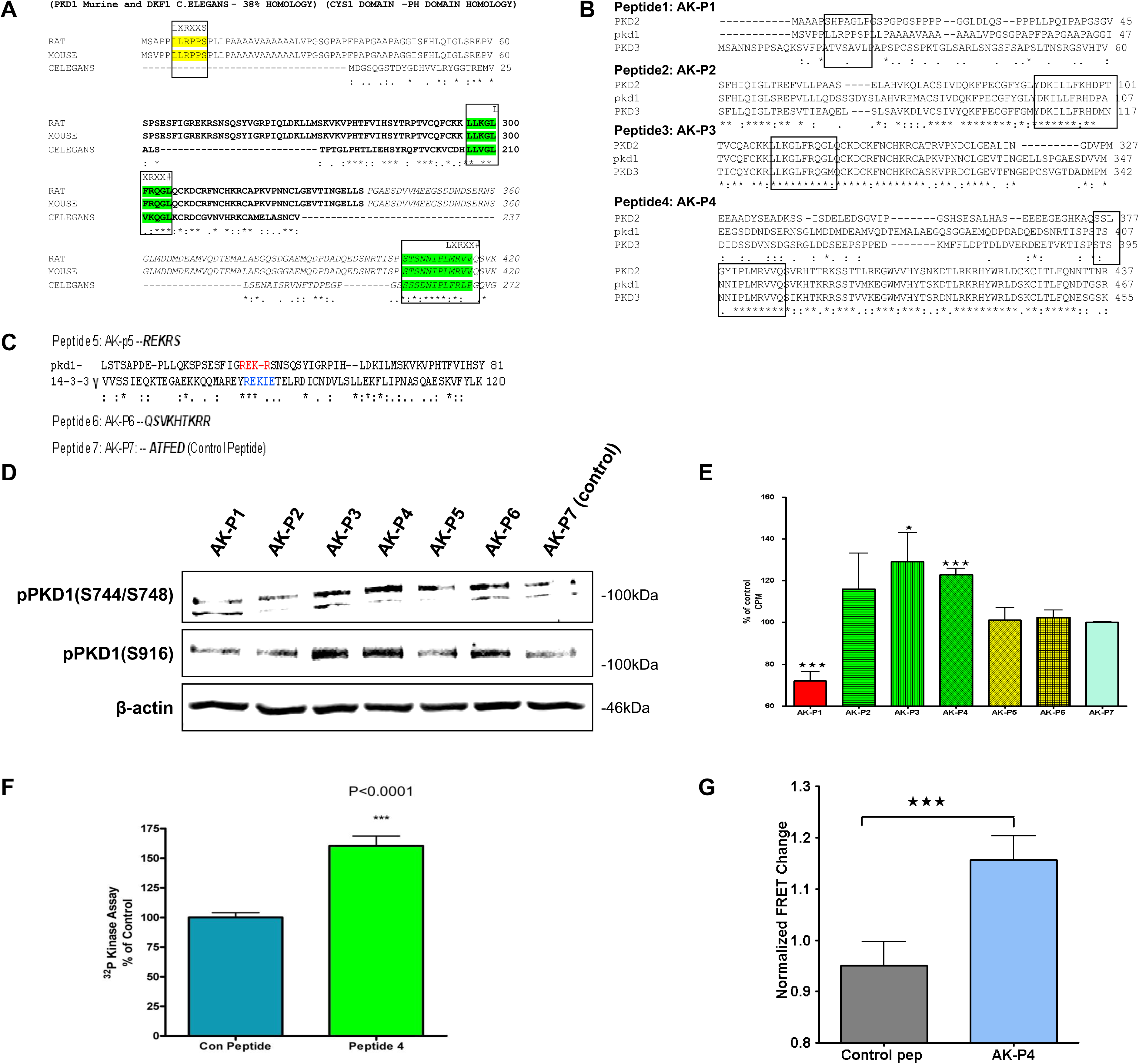
Rationally designed peptide activates PKD1. Alignment of amino acid sequences of PKD1 regulatory fragment between distantly related different organisms (rat, mouse and C. elegans) shows the presence of the consensus substrate-like motif (LXRXXS/T) and pseudosubstrate-like motif (LXRXX#, LXR##, #-serine or threonine residue replaced mostly by a hydrophobic/aliphatic residue) (A). Alignment of mouse PKD isoforms PKD1, PKD2, PKD3 amino acid sequences containing the four (AK-P1, AK-P2, AK-P3, AK-P4) substrate-like (LXRXXS/T) and pseudosubstrate-like region shows high level of conservation (B). 5-aa peptide AK-P5 (54-58 amino acids) from the cysteine rich domain 1 (pseudo-14-3-3γ), a 9-aa peptide AK-P6 (79-87 amino acids) from the pleckstrin homology (PH) domain (pseudo-Gβγ) and a 5-aa scramble control peptide AK-P7 (135-139 amino acids) were synthesized (C). Primary screening: N27 cells were treated with the peptides were probed for PKD1 (pS744E/pS748E) at 60 min (D). Secondary screening: N27 cells were treated with the peptides and PKD1 kinase activity was measured using [^32^P] kinase assay; the bands were quantified for the graph (E). [^32^P] kinase assay was performed using recombinant human PKD1 protein in a cell-free system to test the specific and direct role of AK-P4 peptide (F). N27 cells transfected with 5 μM PKD1 kinase activity reporter (PKD1-KAR) plasmid and 5 μM vector plasmid were treated with AK-P4 and monitored for CFP/YFP FRET change (G). *, p<0.05 and ***, p<0.001 denotes significant difference between the groups.

### PKD1 activation by AK-P4 TAT in cell culture models of PD

The effect of PKD1 peptide activators on cell survival was tested in N27 cells using toxins MPP^+^ and 6-OHDA. Fig. 3A shows that N27 cells treated with 50 µM AK-P3 and AK-P4 both protected against 50 µM 6- OHDA-induced toxicity. But only 50 µM AK-P4 not AK-P3 protected against N27 cells treated with 300 µM MPP^+^ (Fig. 3B). Additional validation of the role of AK-P4 was sought by extending these studies to animal models. To facilitate efficient delivery of AK-P4 into the brain, we tagged the AK-P4 peptide with a 13-aa HIV-TAT transporter peptide through diglycine conjugation. Neuroprotective effect of the TAT-conjugated AK-P4 peptide (AK-P4T) was tested initially in N27 cells and mouse primary mesencephalic neuronal cultures. As shown in Fig. 3C, AK-P4T protected against 100 µM 6-OHDA^-^induced toxicity at a concentration of 100 nM. Further, primary mesencephalic neurons were co-treated with 50 nM AK- P4T to determine its protective effect against 5 µM MPP^+^ treatment for 48 hours. As shown in Fig. 3D, the number of surviving tyrosine hydroxylase (TH)-positive neurons in AK-4T co-treatment group was almost equal to control group, whereas MPP^+^ alone treatment group only showed 30%. TH neurite length in AK-P4T co-treatment group was equal to control group, whereas the TH neurite length was reduced 3-fold in MPP^+^-treated neurons (Fig. 3E). Next, the effects of AK-P4T on the functional efficiency of dopaminergic neurons were studied using H^3^ dopamine uptake assay. As shown in Fig. 3F, AK-P4T co-treatment brought back the dopamine uptake of MPP^+^-treated neurons to 80% of control, whereas MPP^+^ treatment group only showed 50% dopamine uptake. Visualization of TH neuron morphology through fluorescence microscopy shows clear neuroprotective effect of AK- P4T (Fig. 3G). Together, the results demonstrate the protective effect of AK- P4T convincingly in PD cell culture models.

**Figure 3.**
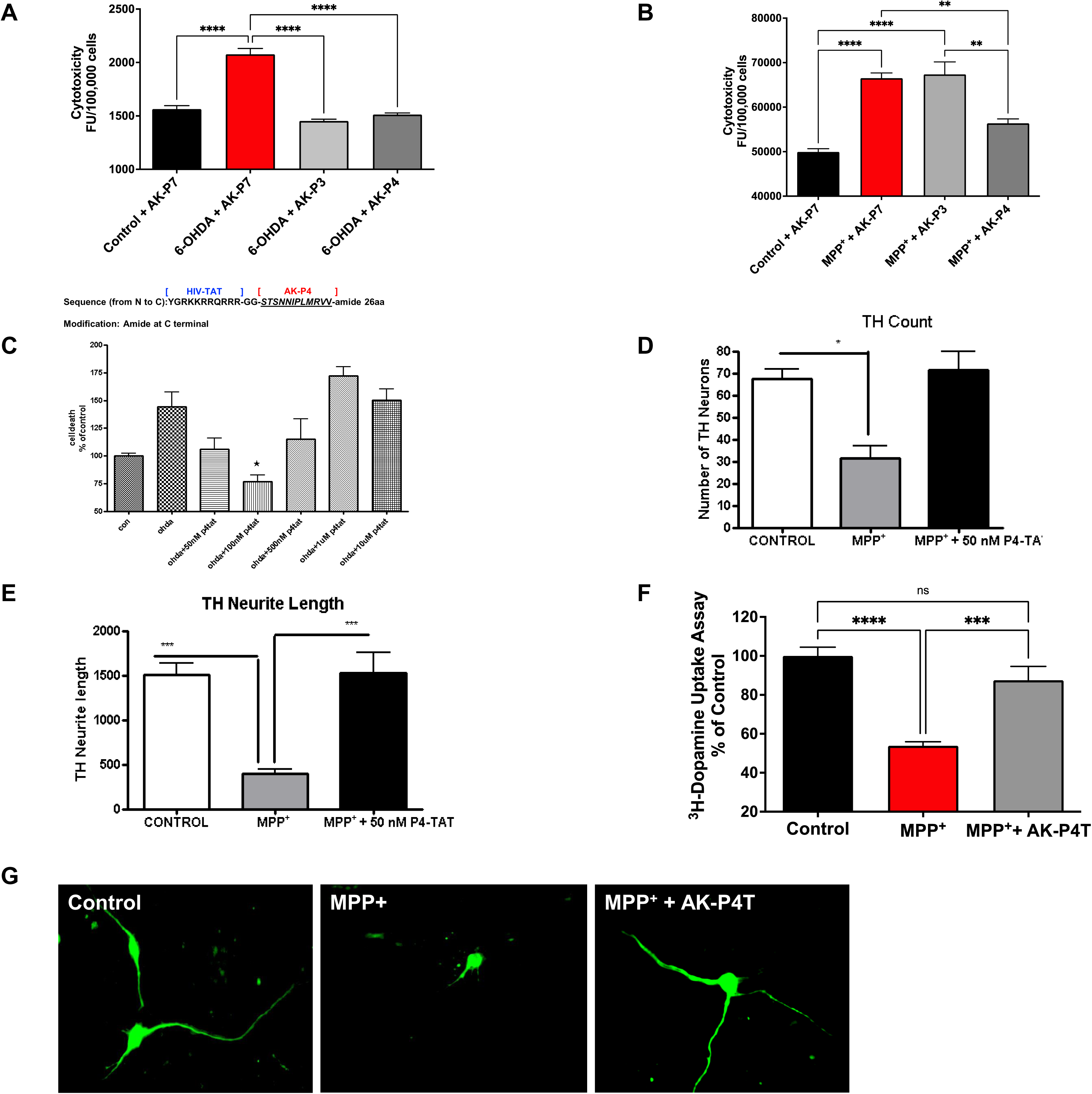
AK-P4 protects against neuronal death in dopaminergic neurons. N27 dopaminergic cells were treated with 6-OHDA (100 μM) and MPP^+^ (300 μM) with or without control peptide (AK-P7), AK-P3, AK-P4 and probed for cytotoxicity (A, B). N27 cells were treated with 6-OHDA (100 μM) and co-treated with different doses (0-10 μM) AK-P4T (AK-P4 was tagged with aa HIV-TAT transporter peptide) and probed for cytotoxicity (C). Mouse primary mesencephalic neurons were co-treated with or without 50 nM AK-P4T and monitored for the number of protected TH neurons (D), TH neurite length (E), neurotransmission function using ^3^H dopamine uptake assay (F), and TH neuronal morphology (G). *, p<0.05, **, p<0.01, and ***, p<0.001 denote significant difference between groups. ns, not significant.

### Intravenous delivery of AK-P4T effectively activates PKD1 in animal models

After characterizing the neuroprotective effect of the activator peptide (AK-P4T) in cell culture models of PD, we examined the ability of AK-P4T to activate PKD1 in the brain. First, we tested different methods to deliver the activator peptide effectively into the brain. Different doses of AK- P4T peptide were delivered intra-peritoneally using different administration paradigms. The intra-peritoneal Ak-P4T delivery of 5 mg/kg activated PKD1 in mice after 4 h of injections but the fold increase of PKD1 in the substantia nigra was not significant as monitored by the activation loop phosphorylation (PKD1pS744/pS748) (Fig. 4A). So, we pursued different routes to deliver AK-P4T. We used nanopolymers to deliver the peptides.

**Figure 4.**
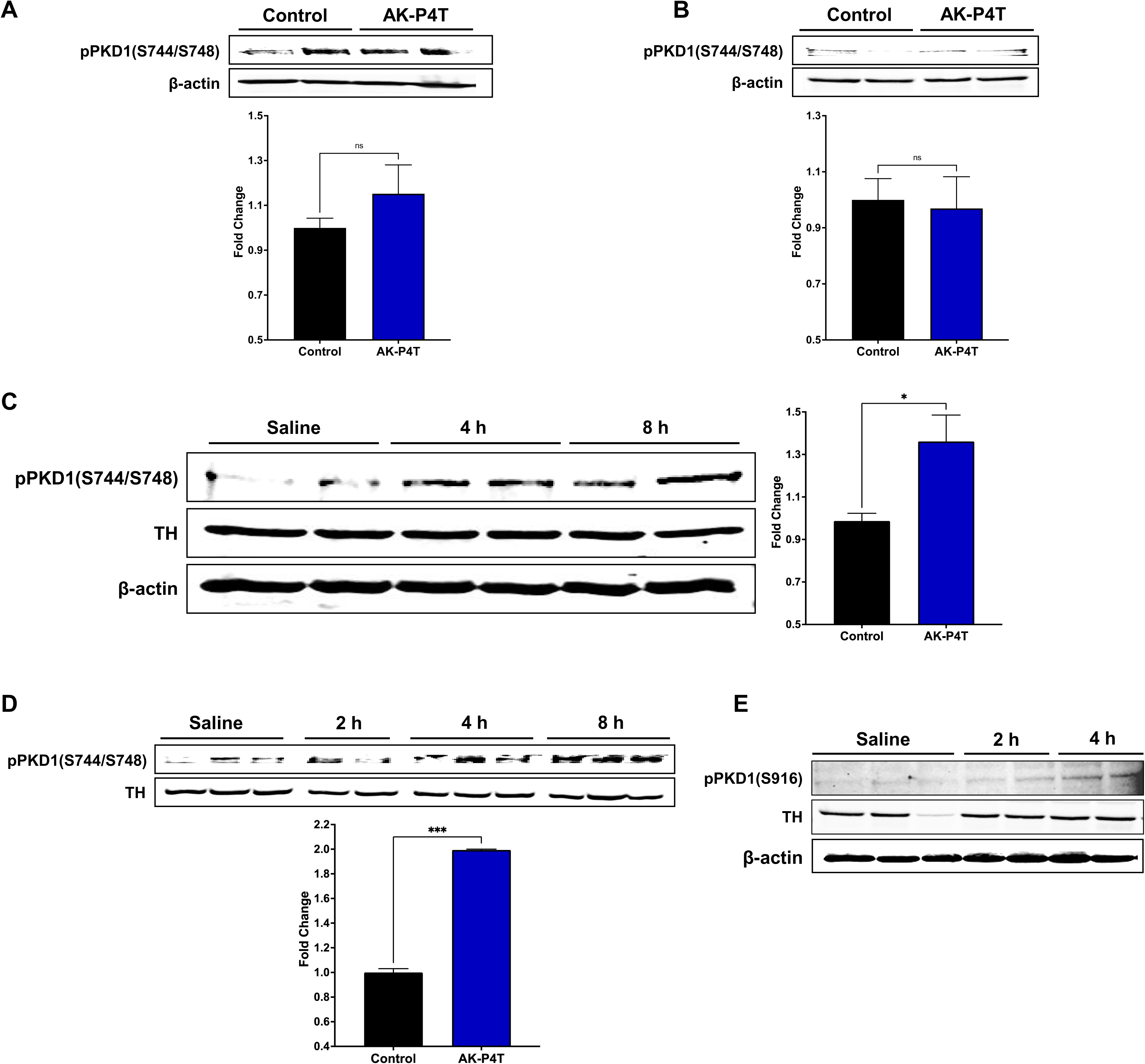
In-vivo delivery of AK-P4T. C57BL/6 mice was administered AK-P4T through intra-peritoneal injections (A), intra-nasal administration of peptides coated with nanopolymers (B), intra-venous administration (C) and monitored for PKD1pS744/pS748 in the substantia nigra (A-C) and striatum (D). C57BL/6 mice was administered AK-P4T through intra-venous injections and monitored for PKD1pS916 in the substantia nigra (E). *, p<0.05 and ***, p<0.001 denotes significant difference between the groups. ns, not significant.

As detailed in the materials and methods section, we used polymer (50:50 CPTEG:CPH) to coat the peptides and delivered them intra-nasally. But the polymer-conjugated AK-P4T peptide through this delivery route in mice did not activate PKD1 in the substantia nigral region (Fig. 4B). So, we tried the intra-venous injections to deliver the peptides. We administered 5 mg/kg peptide through the tail vein of the mice and sacrificed at 4 and 8 hr. As shown in Fig. 4C, the intra-venous AK-P4T peptide delivery activated PKD1 in the substantia-nigra starting at 4 h. We see a similar trend of PKD1 activation in striatum starting at 4 h (Fig. 4D). Quantification of the activation loop phosphorylation (PKD1^pS744/pS748^) at 8 h shows a 1.4-fold increase in PKD1 activation in the substantia nigra and a 2-fold increase in PKD1 activation in the striatum (Fig. 4C and D). Further, intra-venous administration of AK-P4T also increased PKD1^S916^ phosphorylation starting at 4 h confirming the AK-P4T allosteric activation of PKD1 (Fig. 4E). Collectively, these results show that intra-venous delivery of the AK-P4T peptide can activate PKD1 effectively.

### AK-P4T protects against dopaminergic degeneration in an MPTP PD animal model

Next, we examined the neuroprotective efficacy of AK-P4T in a sub- acute MPTP mouse model of PD. Since AK-P4T activated PKD1 through intra-venous administration, we chose this delivery route. Animals were administered AK-P4T (5 mg/kg body weight) 24 h before MPTP administration. Animals were then cotreated with AK-P4T and MPTP (25 mg/kg i.p.) daily for 5 days and the AK-P4T treatment was continued for 7 more days and the animals were sacrificed. Saline- and vehicle-injected animals were used as controls. To assess the protective effect of AK-P4T, we first examined whether AK-P4T could block MPTP-induced loss of striatal dopamine and its metabolites. As shown in Fig. 5A, MPTP treatment induced loss of dopamine (>75%) in mouse striatum. Pretreatment with 5 mg/kg AK-P4T afforded more than 50% protection against MPTP-induced striatal dopamine loss. The dopamine levels were determined to be 72.21 ± 16.67, 18.56 ± 2.18 and 38.95 ± 5.365 pg/mg protein in control-, MPTP-, and MPTP + 5 mg/kg AK-P4T-treated animals, respectively (Fig. 5A). Similar results were found for dopamine metabolites DOPAC and HVA (Fig. 5B and C). Further, Western blotting of striatal lysates for TH shows rescued TH levels in the AK-P4T co-treatment group (Fig. 5D). Further, immunochemical DAB staining shows the rescue of TH levels in AK-P4T co-treatment group compared to MPTP group in both striatal and substantia nigra sections (Fig. 5E-F). Western blotting of substantia nigra lysates shows that PKD1 activation is preserved in AK-P4T co-treated groups (Fig. 5G). Collectively, these data suggest that AK-P4T treatment could afford protection against MPTP-induced nigro-striatal dopamine and dopamine metabolite loss in animal PD models by persistent activation of PKD1.

**Figure 5.**
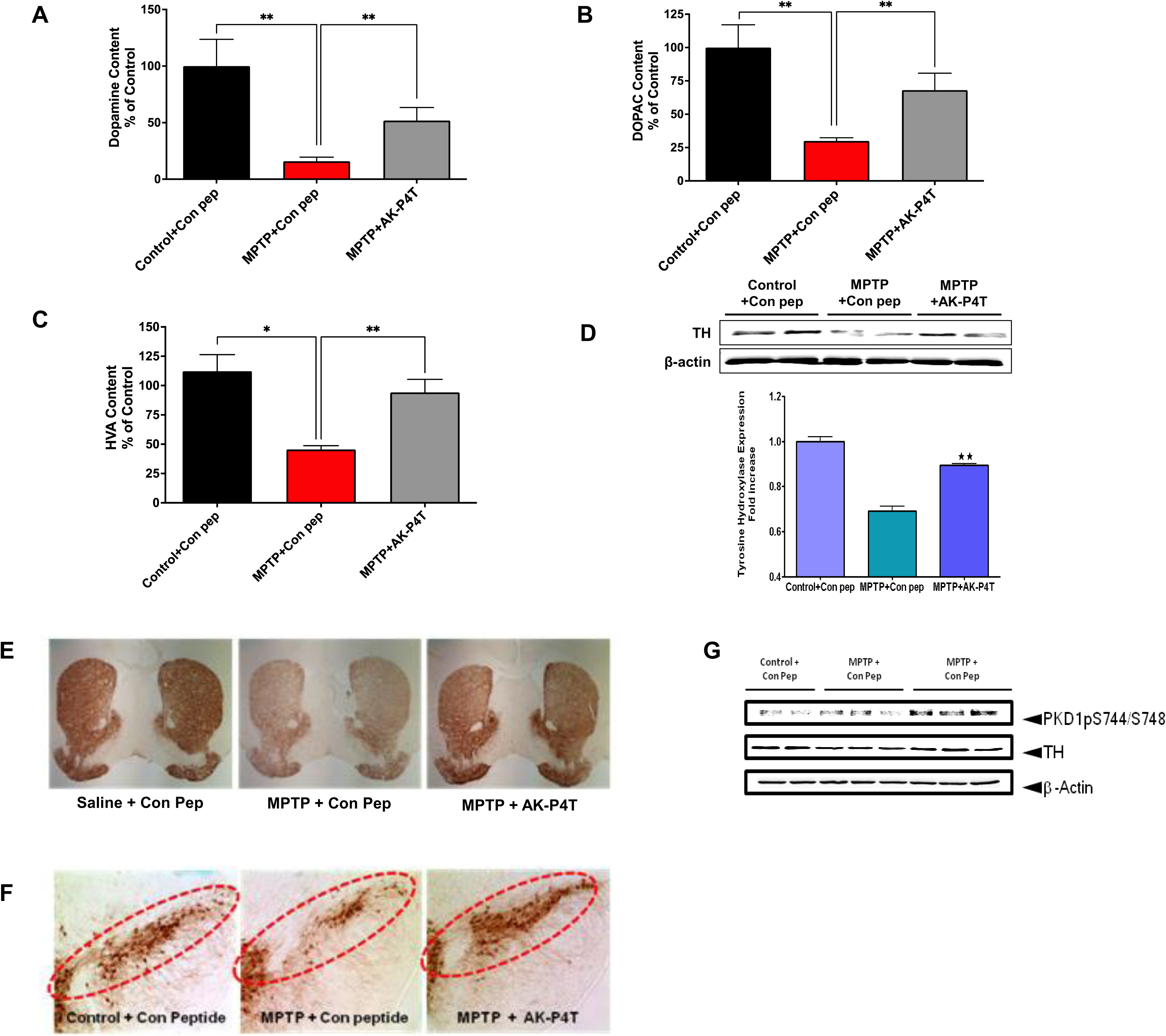
AK-P4T mediated PKD1 activation rescues neurotransmitter levels and dopaminergic function in pre-clinical model of PD. Neuroprotective efficacy of AK-P4T was examined in a sub-acute MPTP mouse model of PD. Animals pre-treated and co-treated with AK-P4T and control peptide (5 mg/kg body weight) were sacrificed at the end of the study and the following analyses were performed: Striatal brain lysates were analyzed for Dopamine (A), DOPAC (B), HVA (C) levels were measured using HPLC. Striatal lysates were blotted for Tyrosine Hydroxylase (TH) (D). Immunohistochemical analysis of dopaminergic neurons staining for tyrosine hydroxylase (TH) DAB staining were performed from Mouse brain slices cut at the level of the striatum (E) and substantia nigra (F). Substantia nigral lysates were blotted for Tyrosine Hydroxylase (TH) and PKD1 pS744/S748 (G). *, p<0.05 and **, p<0.01 denotes significant difference between the groups.

### AK-P4T attenuates motor deficits

With evidence suggesting that AK-P4T protects against MPTP-induced dopamine and TH neuronal loss, we next determined whether AK- P4T treatment could also afford protection against MPTP-induced motor deficits. We compared the motor activity of animals treated with vehicle plus control peptide, MPTP plus control peptide, and MPTP plus AK-P4T peptide, using a VersaMax computerized activity monitoring system (Accuscan, Columbus, OH). This system uses infrared sensors to measure repetitive movements both in the horizontal and vertical planes in real time, and it provides color-coded output. Representative activity maps of the animals are presented in Fig. 6A. The cumulative behavioral activities are monitored and expressed as percentage of the vehicle-treated group, and they were obtained at day 9 of the sub-acute MPTP treatment paradigm. Statistical analyses of both raw data and percentage of control showed significant differences in various behavioral activities. The data indicates significantly reduced horizontal motor activity (>20% reduction), number of movements (>20% reduction), vertical movement time (>50% reduction), vertical motor activity (>40% reduction), total distance travelled (>20% reduction), vertical number of movements (>40% reduction), movement time (>20% reduction), stereotypy count (>20% reduction), margin distance (>20% reduction), margin time (>20% reduction), stereotypy time (>20% reduction), rearing activity (>40% reduction), and rearing time (>50% reduction) in MPTP-treated animals compared with the vehicle-treated group (Fig. 6B-N). However, administration of AK-P4T almost completely restored all the behavioral and locomotory patterns of MPTP-treated animals to the levels observed in control animals (Fig. 6B-N). These results demonstrate AK-P4T treatment attenuates MPTP-induced locomotor deficits in a pre-clinical model of PD.

**Figure 6.**
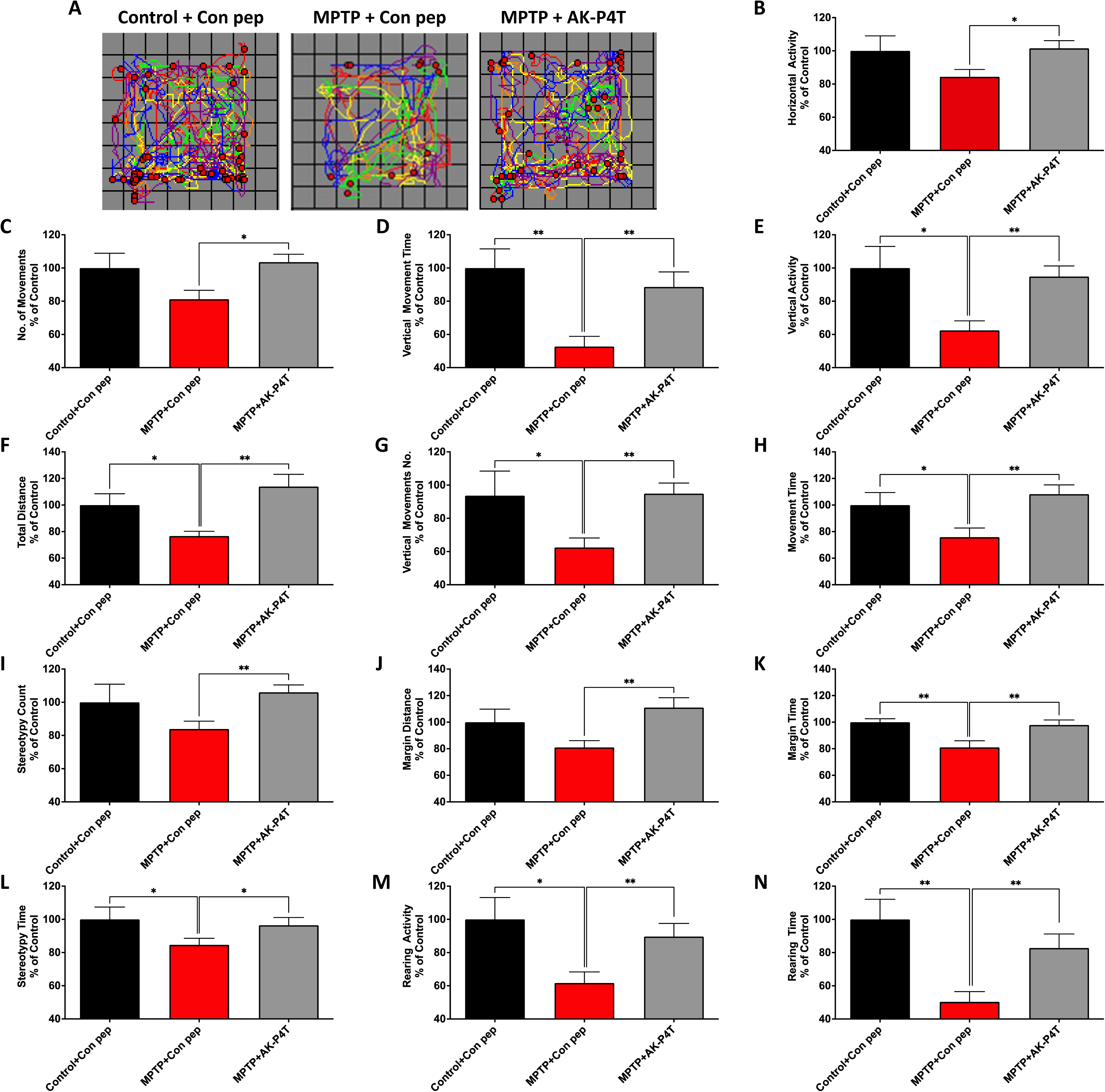
AK-P4T completely attenuates behavioral and locomotory deficits in a pre-clinical PD model. Behavioral and locomotory changes were measured after AK-P4T and control peptide (5 mg/kg body weight) co-treatment in the sub-chronic MPTP mouse model of PD. (A) Representative activity maps of the animals are shown. (B-N) The following parameters were measured using VersaMax: horizontal motor activity, number of movements, vertical movement time, vertical motor activity, total distance travelled, vertical number of movements, movement time, stereotypy count, margin distance, margin time, stereotypy time, rearing activity, and rearing time. *, p<0.05 and **, p<0.01 denotes significant difference between the groups.

## Discussion

The present study demonstrates that a rationally designed allosteric activator peptide of PKD1 protects against dopaminergic neurodegeneration in pre-clinical models of Parkinson’s disease. Extensive characterization of PKD1 signal transduction in pre-clinical models of PD showed that, positive modulation of PKD1 can be a novel therapeutic strategy against PD. Herein, we report for the first time some key findings pertinent to neuroprotection against Parkinson’s disease: (i) rationally designed peptide modulators based on the substrate-like and pseudo- substrate like sequences present in the regulatory fragment activate or inhibit PKD1 allosterically by mimicking and disrupting the native intra-protein interactions, (ii) one of the peptide activators AK-P4 protected against both 6-OHDA- and MPP^+^-induced toxicity in N27 dopaminergic cells, (iii) TAT_47-57_ peptide-conjugated AK-P4 (AK-P4T) rescued TH^+^ neurons against MPP^+^ and prevented dopaminergic neurodegeneration in primary mesencephalic cultures, and (iv) effective delivery of AK-P4T through intravenous administration activated PKD1 and protected against MPTP-induced motor deficits, striatal dopamine depletion, and nigral dopaminergic neuronal loss in a sub-acute MPTP animal model. To our knowledge, this is the first demonstration of a novel neuroprotective strategy against PD by positively modulating an anti-apoptotic kinase PKD1 using a rationally designed allosteric activator.

Based on these findings we hypothesized that prolonging the activation of the survival switch PKD1 through a specific activator can prevent or delay PD progression. Recently, peptide-based drugs have successfully entered clinical trials or approved as a treatment for many disorders [33, 34]. Additionally, they are specific with fewer side effects compared to small molecule drugs [35]. We utilized the vast protein databases and developed a new approach to find PKD1 modulators. We began with our hypothesis that interfering with intra-protein interactions of PKD1 can modulate PKD1. PKD1 in native state is auto-inhibited by its regulatory fragment consisting of the cystine rich domains and pleckstin homology (PH) domain but do not have a pseudo-substrate region like some PKC family members. Careful analysis of PKD1 protein sequence revealed highly conserved regions similar to the consensus substrate sequence. Our results suggest that peptides synthesized mimicking the pseudo-substrate like region activates the kinase, while peptide mimicking the substrate like region inhibits the kinase. Particularly, the peptide AK-P4 mimicking the highly conserved pseudo-substrate like region in the PH domain protected against both 6-OHDA and MPP^+^ mediated toxicity.

One of the disadvantages for peptide-based therapeutics is rapid drug clearance derived from inherent properties of peptides. In gastrointestinal system, they can be easily hydrolyzed by digestive enzymes [33]. Several studies have proved abilities of cell penetrating peptides (CPP), such as TAT or TP10, to penetrate cell membrane and deliver protein to targets [36–38]. Among other types of CPPs, CPP-derived the trans-activating transcriptional activator (TAT) from HIV can be used to deliver therapeutic peptides [39, 40]. A study showed that ND-13, designed a short peptide, is composed of a DJ-1 segment attached to 7 amino acids derived from TAT [41]. The peptide not only protected cells from oxidative stress in vitro but also restored dopaminergic functions both in 6-OHDA, MPTP, and DJ-1 knockout mouse models.

AK-P4 peptide was conjugated with the TAT_47-57_ (AK-P4T) to facilitate effective delivery across the blood-brain barrier. Even though, progress has been made to deliver therapeutic peptides for neurodegenerative disorders, effective peptide drug delivery remains as an area for improvement [42]. Recently, some research has proved that drug administration by intra-nasal route is convenient and effective because it directly targets to brain via olfactory neurons [43, 44]. However, one of limitations is nasal mucociliary clearance, unavailable to stay longer time in nasal cavity. Therefore, intra-venous administration can be a reasonable option without gastrointestinal absorption even if it is not convenient for everyday use [44]. We tested different routes and methods to effectively deliver the peptide; inta-peritoneal administration, intra-nasal administration of peptides coated with nanopolymers, intra-venous administration. Our results indicate that only AK-P4T peptide administered through intra-venous route can activate PKD1 in the nigro-striatal pathway after effectively crossing the blood-brain barrier. We used intra-venous routes for all our studies, even though exploration of other methods is underway.

Our findings in pre-clinical models using MPTP demonstrate that AK- P4T delivered through intra-venous route activates PKD1 and rescues dopaminergic neurons from succumbing to cell death. Further, the nigro- striatal dopaminergic neuronal loss is reduced improving the neuro- transmission function of dopaminergic neurons. This considerably improves the motor function as the mice behave like the control group.

In conclusion, we demonstrate that activation of the novel protective signaling mechanism mediated by PKD1 in dopaminergic neurons by a peptide-based PKD1 activator offers neuroprotection in pre-clinical models of PD by slowing down or stopping dopaminergic degeneration (Fig. 7). This drug offers therapeutic promise for PD.

**Figure 7.**
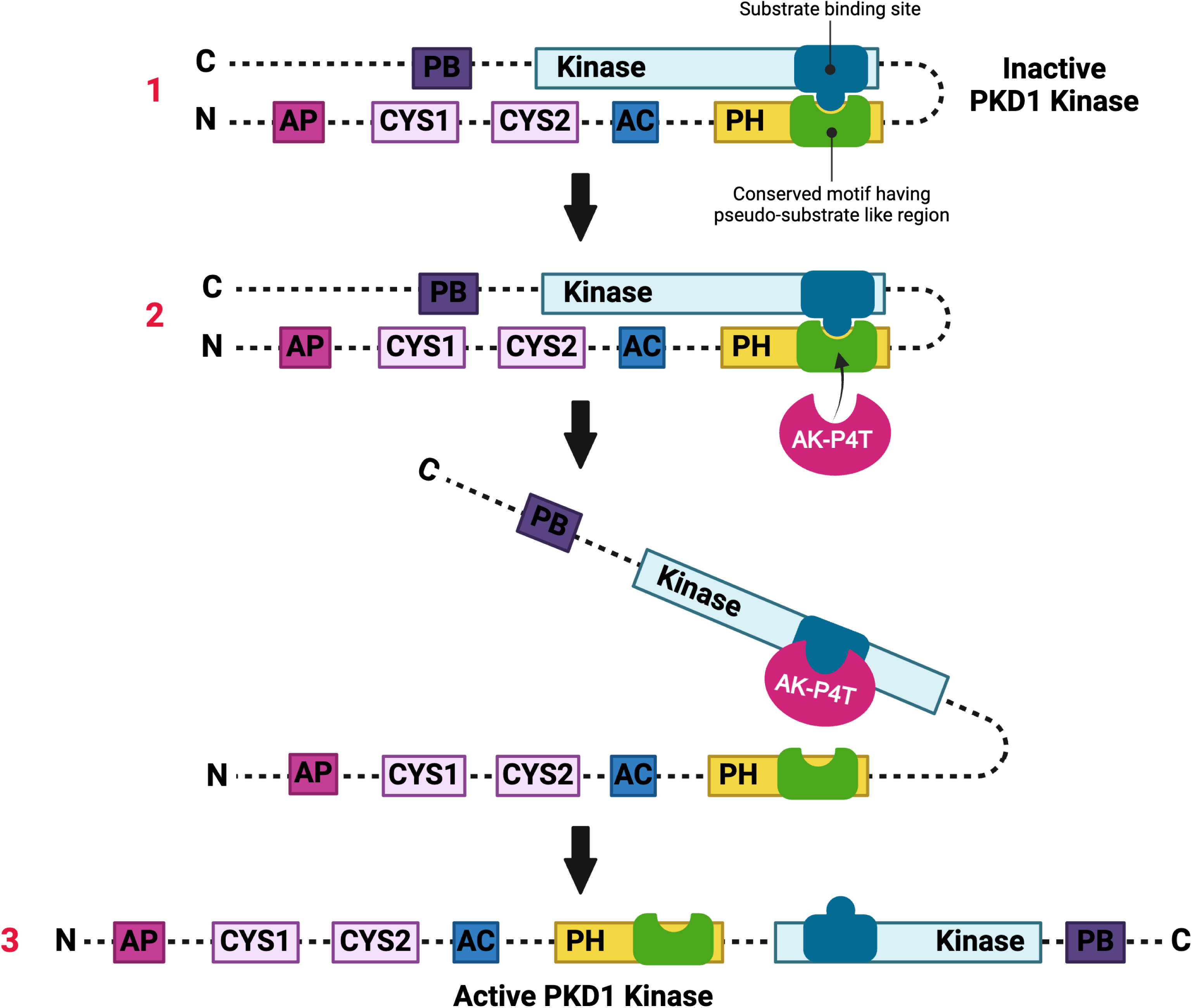
A working model for AK-P4T activation mechanism. 1. The regulatory fragment of PKD1 auto-inhibits its catalytic fragment through inhibitory pockets containing the pseudo-substrate-like sequences. 2. Peptide regulators (AK-Series) mimic the inhibitory pockets, interact with the catalytic fragment and release the auto-inhibition. 3. Persistent activation of PKD1 promotes neuronal survival.

## Abbreviations

PD: Parkinson’s disease
PKD1: Protein kinase D1
MAPK: Mitogen-activated protein kinases
PKCδ: Protein kinase C delta
CAMK: Ca^2+^/Calmodulin-Dependent Protein Kinase II
JNK: c-Jun N-terminal kinases
LRRK2: Leucine-rich repeat kinase 2 (LRRK2)
MLK: Mixed-lineage kinase
ROS: Reactive oxygen species
MnSOD: Manganese superoxide dismutase
WB: Western Blot
PKC: Protein kinase C
PKCα: Protein kinase C alpha.

## Authors’ contributions

**Arunkumar Asaithambi**: Conceptualization, Investigation, Methodology, Data curation, Visualization, Validation, Formal Analysis, Writing—original draft, Writing—review and editing. **Ahyoung Jang**: Formal Analysis, Writing-review and editing. **Anamitra Ghosh**: *Investigation, Data curation, Methodology, Validation*. **Muhammet Ay**: *Investigation, Data curation, Methodology, Validation*. **Huajun Jin**: *Writing—review and editing*. **Vellareddy Anantharam**: *Project Administration, Resources, Writing— review and editing*. **Arthi Kanthasamy**: *Conceptualization, Funding Acquisition, Resources*. **Anumantha Kanthasamy**: *Conceptualization, Funding Acquisition, Project Administration, Resources, Supervision, Writing—review and editing*.

## Acknowledgements

The authors acknowledge Mr. Gary Zenitsky for his assistance in the preparation of this manuscript. This work was funded by the National Institute of Health grants NS121692, ES034196 and NS121692. Other sources include the Johnny Isakson Endowment the Coach Mark Richt Neurological Disease Research Fund to AGK and AK.

